# Does the ratio of β-1,4-glucosidase (BG) to β-1,4-N-acetylglucosaminidase (NAG) indicate the relative resource allocation of soil microbes to C and N acquisition?

**DOI:** 10.1101/2021.02.08.430361

**Authors:** Taiki Mori, Ryota Aoyagi, Kanehiro Kitayama, Jiangming Mo

## Abstract

The ratio of β-1,4-glucosidase (BG) to β-1,4-N-acetylglucosaminidase (NAG) activity (BG:NAG ratio) is often used as an indicator of the relative resource allocation of soil microbes to C acquisition compared with N. An increasing number of recent studies have used this index to assess the nutrient status of microbes. However, the validity of this index for assessing the nutrient status of microbes is not well tested. In this study, we collected published data and tested that validity by investigating whether N fertilization elevated the BG:NAG ratio, assuming that microbes reduce their allocation to the N-acquiring enzyme (NAG) under N-enriched conditions. Of the data points, 54% (82/151) did not support the hypothesis because those studies showed lower BG:NAG ratios in N-enriched soils than under ambient conditions, especially when the ambient BG:NAG ratio was higher than 2.0 (77%, 59/77). This suggests that the BG:NAG ratio does not always indicate the microbial status for C or N limitation. Rather, we hypothesized that the decomposition stage explained the variation in BG:NAG because N addition accelerates decomposition, and the BG:NAG ratio is lower at later stages of decomposition due to the dominance of NAG-targeting C (chitin or peptidoglycan). A negative correlation of BG:NAG ratio with polyphenol oxidase activity, which increases with decomposition, supported our hypothesis.

## Introduction

Prior authors have suggested that the relative resource allocation of soil microbes to acquire energy and nutrients can be expressed as the ratio of extracellular enzyme activity targeting carbon (C), nitrogen (N), and phosphorus (P) (Sinsabaugh *et al*., 2008, 2009; Waring *et al*., 2014). The ratio of β-1,4-glucosidase (BG) to β-1,4-N-acetylglucosaminidase (NAG) (BG:NAG ratio) is often used as an indicator of the relative resource allocation of microbes to C acquisition compared with N acquisition (Turner & Wright, 2014; Waring *et al*., 2014; Zhou *et al*., 2017; Chen *et al*., 2018). Recently, increasing numbers of studies have used the BG:NAG ratio to assess nutrient limitation or the status of microbes, assuming that a lower BG:NAG ratio indicates N shortage/N limitation (hereafter, the ecoenzymatic stoichiometry hypothesis; (Sinsabaugh *et al*., 2008; Waring *et al*., 2014; Moorhead *et al*., 2016; Chen *et al*., 2018; Mori *et al*., 2018a; Wang *et al*., 2018)). Nevertheless, this hypothesis has not been tested sufficiently, and several studies examining it have reported inconsistent results (Mori *et al*., 2018a; Rosinger *et al*., 2019; Mori, 2020). The validity of the BG:NAG ratio as a measure of the nutrient status of microbes needs to be tested by synthesizing accumulated data.

The ecoenzymatic stoichiometry hypothesis can be tested in N fertilization experiments. If the BG:NAG ratio really indicates the nutrient status of microbes, N fertilization will elevate it because microbes can get N directly from the fertilizer and reduce the allocation to N-acquiring enzymes, *i.e.,* NAG. Furthermore, the response of the BG:NAG ratio to N fertilization would be smaller under N-rich conditions because the decrease in NAG activity would be smaller. Accordingly, the predicted distribution of data could be drawn as shown in Fig. 1. In the present study, we collected published papers reporting the impact of N fertilization on the activity of BG and NAG and compared the BG:NAG ratio in surface soils before and after N fertilization.

**Fig. 1.**
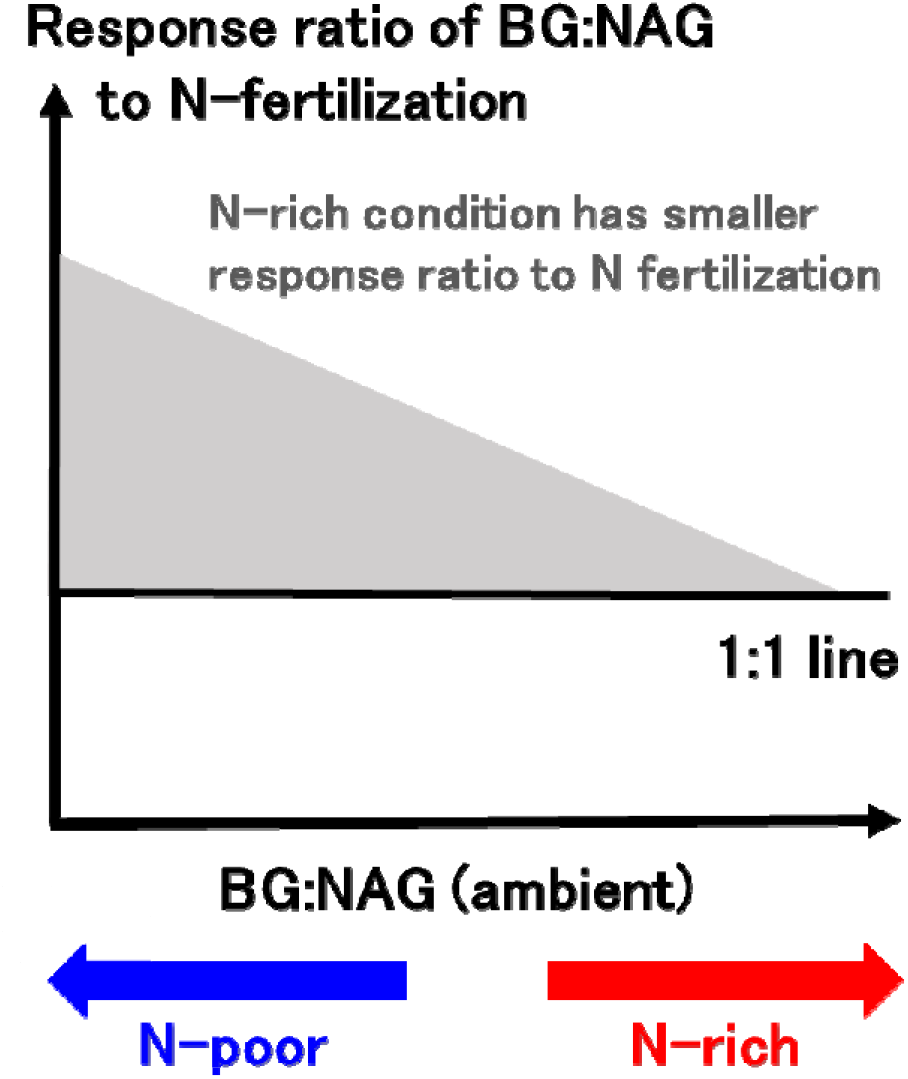
The predicted relationship between the BG:NAG ratio under ambient conditions and the response ratio of the BG:NAG ratio to N fertilization according to the enzymatic stoichiometry hypothesis. Data points would be plotted above the 1:1 line because N fertilization does not lower the BG:NAG ratio. N-rich conditions (*i.e.,* a low BG:NAG ratio) showed a lower response ratio to N fertilization.

We also established an alternative hypothesis, which stems from a completely different mechanism: the BG:NAG ratio does not indicate the relative allocation of microbes to C and N, but the source of the C resources (substrate) for microbes (Mori, 2020). This hypothesis is plausible for the following two reasons: (i) enzyme activity varies depending on the relative substrate availability [several studies have reported that the addition of a substrate caused elevation of enzyme activity that targets the added substrate (Shackle *et al*., 2000)], and (ii) both BG and NAG can be produced for acquiring C (Mori *et al*., 2018a; Wang *et al*., 2018; Mori, 2020). When cellulose (*i.e.,* a BG-targeting C compound) is the dominant C resource in soil, microbes utilize more cellulose than chitin and peptidoglycan (a NAG-targeting C compound), leading to higher BG activity and a higher BG:NAG ratio (Mori, 2020). By contrast, chitin and peptidoglycan-dominant conditions cause microbes to utilize more chitin and peptidoglycan than cellulose, resulting in higher NAG activity and a lower BG:NAG ratio (Mori, 2020). As decomposition progresses, the relative abundance of cellulose decreases, whereas chitin and peptidoglycan becomes more abundant because chitin and peptidoglycan derived from microbial death is supplied to the soil via microbial turnover (see Fig. 2). As a result, NAG becomes a more dominant C-acquiring enzyme than BG. Therefore, our alternative hypothesis predicts that the BG:NAG ratio is lower when abundant soil organic matter is progressively decomposed (Fig. 2). According to this hypothesis, N fertilization can reduce the BG:NAG ratio, in contrast to the enzymatic stoichiometry hypothesis, because N enrichment is expected to stimulate organic matter decomposition under N-poor conditions (Hobbie, 2005). Furthermore, the hypothesis predicts that the BG:NAG ratio will be negatively correlated with the activity of polyphenol oxidase (PPO), a well-measured ecoenzyme that oxidizes lignin or humus and increases as decomposition progresses (Moorhead & Sinsabaugh, 2006; Sinsabaugh & Shah, 2011). To validate the prediction, data on PPO activity were also collected from the literature. As N fertilization often has negative effects on PPO, we analyzed the correlations using only N-unfertilized data.

**Fig. 2.**
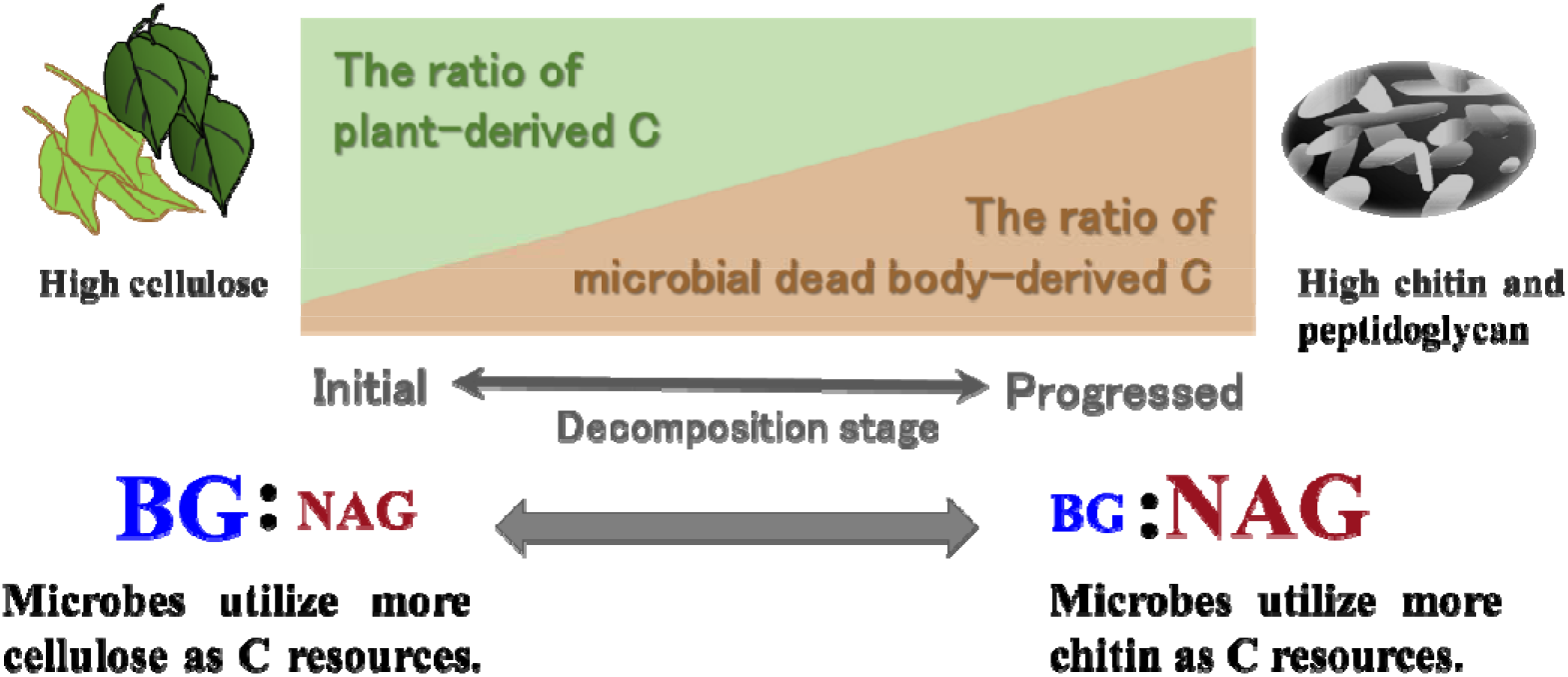
Our new hypothesis explaining what the BG:NAG ratio shows. As microbial activity accelerates, the ratio of plant-derived C in the total C pool decreases, while the ratio of microbial dead body-derived C in the total C pool increases. As a result, NAG becomes the more dominant C-acquiring enzyme compared with BG, which leads to a decrease in the BG:NAG ratio.

## Material and Methods

Jian *et al*. (2016) published a meta-analysis reporting the impact of N fertilization on ecoenzymes, which comprehensively collected data on the response of ecoenzyme activity, including BG and NAG, to N fertilization through 2015, so we used the reported data in our analysis. Then, we searched the Web of Science for papers published later than 2015 using the following combinations of key words: (NAG OR chitinase OR β-1,4-N-acetyl-β-glucosaminidase OR “N-acetyl β-glucosaminidase” OR glucosaminidase) *AND* (BG OR β G OR β-1,4-glucosidase OR glucosidase) *AND* (“nitrogen add*” OR “N add*” OR “nitrogen elevat*” OR “N elevat*” OR “nitrogen fertiliz*” OR “N fertiliz*” OR “nitrogen appl*” OR “N appl*”OR “nitrogen enrich*” OR “N enrich*”) (time span 2015–2018). We collected 151 data points from 40 papers. The relationship between the BG:NAG ratio under ambient conditions and that under N-enriched (fertilized) conditions was examined. If the paper reported PPO activity, we also recorded that. Pearson’s test was used to test the correlations between enzyme activities and the BG:NAG ratio. All statistical analyses were performed using R ver. 3.4.1 or 3.4.4 (R Core Team, 2018).

## Results and Discussion

A large proportion of the synthesized data did not support the ecoenzymatic stoichiometry hypothesis, although the distribution of the data was close to the predicted pattern (Fig. 1) when the ambient BG:NAG ratio was low (Fig. 3b). Of 151 data points, 82 (54%) indicated lower BG:NAG ratios in N-enriched soils than under ambient conditions (*i.e*., the data points are below the solid line in Fig. 3), especially when the ambient BG:NAG ratio was higher than 2.0 (77%, 59/77). According to the chi-square test, a higher positive response ratio occurred when the ambient BG:NAG ratio was <2.0 (*P* < 0.001). The ecoenzymatic stoichiometry hypothesis cannot explain the lowered BG:NAG ratio in N-fertilized soils because, according to that hypothesis, the result indicates that N fertilization enhances the N shortage, which is contradictory. Our meta-analysis suggested that the BG:NAG ratio does not always indicate the nutrient status of soil microbes.

**Fig. 3.**
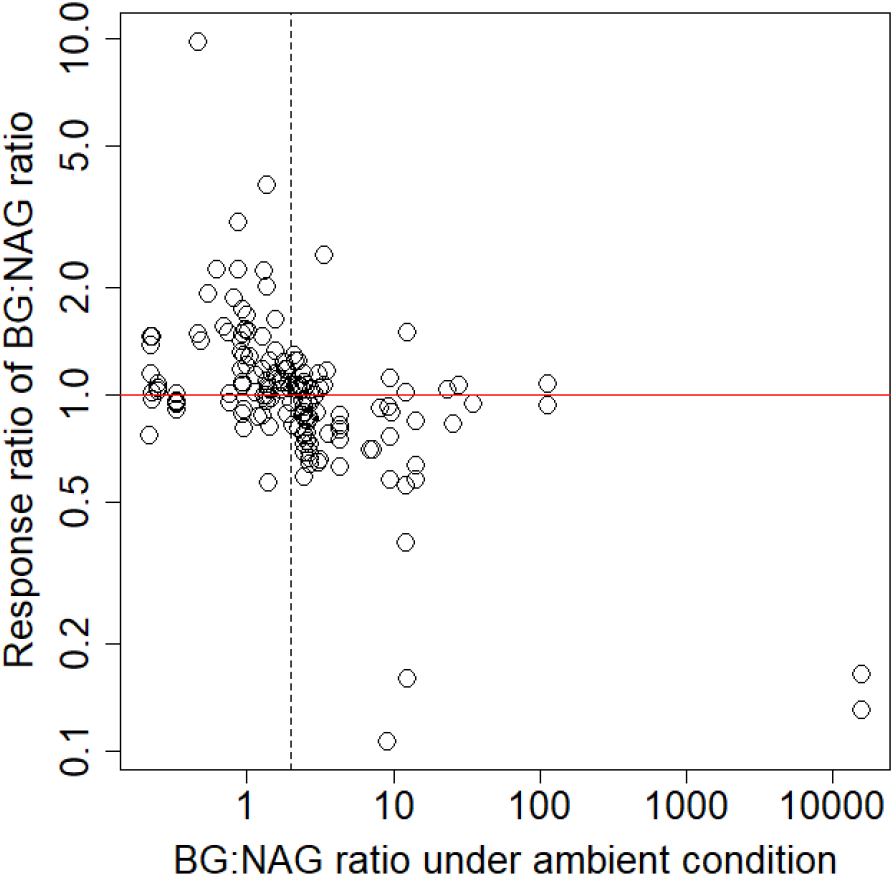
The correlation between the BG:NAG ratio under ambient conditions with the response of the BG:NAG ratio to N fertilization. The solid red line is the 1:1 line. The dashed line represents the ambient condition under which the BG:NAG is 2.0. Of 151 data points, 82 (54%) showed lower BG:NAG ratios under ambient conditions. When the ambient BG:NAG ratio was higher than 2.0, 59 of 77 (77%) data points showed a lower BG:NAG ratio under ambient conditions.

Other studies have also reported results inconsistent with the ecoenzymatic stoichiometry hypothesis (also see a perspective by Mori, 2020). Waring *et al*. (2014) collected 17 studies of N-rich tropical ecosystems and found that the mean BG:NAG ratio in these sites was not significantly different from the global average. In a lowland tropical rainforest in Bornean Malaysia, Mori *et al*. (2018a) reported that the BG:NAG ratio was similar to (or even slightly smaller than) the global average, although the forest is considered N rich (Aoyagi & Kitayama, 2016). This could be explained by the following hypothetical mechanism: under N-rich conditions, microbes produce NAG-targeting C because chitin and peptidoglycan, whose terminal reaction is catalyzed by NAG (Waring *et al*., 2014), contains both C and N (Mori *et al*., 2018a, 2018b; Wang *et al*., 2018). Several papers have supported our idea, reporting high NAG activity with a lack of a response of the NAG activity or BG:NAG ratio to N fertilization (see the meta-analysis by Jian *et al*. 2016).

Our alternative hypothesis was supported by the collected data. As predicted by the hypothesis, PPO activity was negatively correlated with the ambient BG:NAG ratio in our dataset (Fig. 4). The result supports the idea that the BG:NAG ratio shows the decomposition stage. That is, as decomposition progresses, the BG:NAG ratio decreases because the abundance of chitin and peptidoglycan becomes dominant relative to that of cellulose (Fig. 2). Our alternative hypothesis can explain how N fertilization decreases the BG:NAG ratio and why lower BG:NAG ratios were mainly observed when the ambient BG:NAG ratios were >2.0 (Fig. 3). At a relatively early stage of the decomposition process, where the cellulose content is relatively large and dominates the microbial C resources (higher BG:NAG ratio), N fertilization can accelerate decomposition and microbial activity (Yoshitake *et al*., 2007). In such a case, the added N can change the C source of microbes to more microbial dead body-derived C (chitin and peptidoglycan dominant, NAG-targeting C), resulting in a lower BG:NAG ratio under N-added conditions. Conversely, at a later stage of decomposition, N addition generally suppresses organic matter decomposition (Fog, 1988; Knorr *et al*., 2005; Janssens *et al*., 2010), and a lower BG:NAG ratio caused by N fertilization should be observed less often (Fig. 3).

**Fig. 4.**
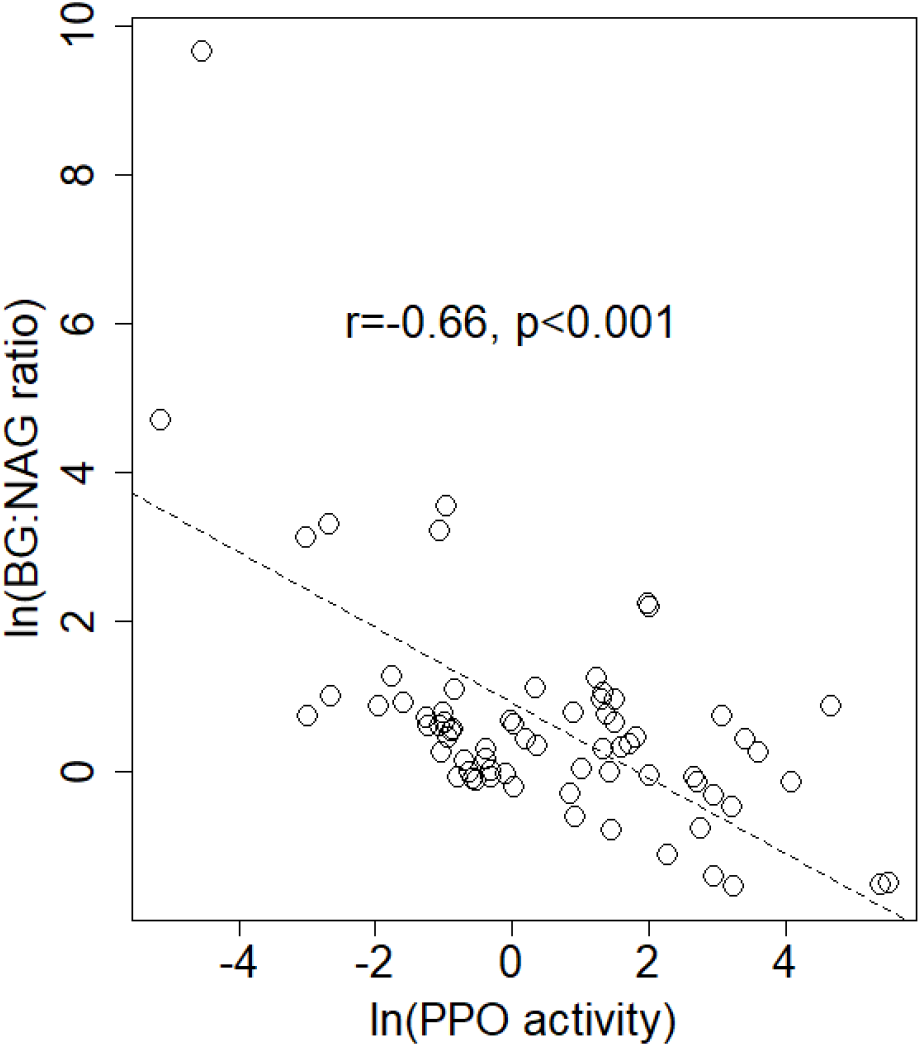
The correlation of ln(PPO activity under ambient conditions) with ln(BG:NAG ratio under ambient conditions).

Although some of our data did not support the ecoenzymatic stoichiometry hypothesis, we cannot completely reject the hypothesis. When the ambient BG:NAG ratios are <2.0, the distribution of the data was close to the pattern predicted by that hypothesis (Fig. 1), and 69% (51/74) of the data points showed higher BG:NAG ratios in N-fertilized soils than under ambient conditions (Fig. 3). It is possible that the ecoenzymatic stoichiometry hypothesis is true when the NAG activity is relatively high, whereas another mechanism (such as our alternative hypothesis) controls the BG:NAG ratio in ecosystems with high BG:NAG ratios. The meta-analysis approach is unable to test this idea. We need other approaches, such as a laboratory experiment monitoring the BG:NAG ratio of organic matter over the course of decomposition under both manipulated N-shortage and N-rich conditions.

In summary, we demonstrated that the BG:NAG ratio may not always indicate the nutrient status of microbes, as previously suggested (Rosinger *et al*., 2019; Mori, 2020), at least when the initial BG:NAG ratio exceeds 2.0. Our dataset also indicated that the stage of decomposition can explain variation in BG:NAG.

## Acknowledgement

This study was financially supported by a grant from environmental research aid, The Sumitomo Foundation 153082 and Grant-in-Aid for JSPS Postdoctoral Fellowships for Research Abroad (28·601) to T Mori.

## Statement of authorship

TM conceived this study. TM and RA wrote the first draft of the manuscript. All of the authors contributed to the discussion and writing of the manuscript.

